# 3D Bioprinting of Architected Hydrogels from Entangled Microstrands

**DOI:** 10.1101/2019.12.13.870220

**Authors:** B. Kessel, M. Lee, A. Bonato, Y. Tinguely, E. Tosoratti, M. Zenobi-Wong

**Affiliations:** Department of Health Sciences and Technology, ETH Zurich, Switzerland

## Abstract

Hydrogels are an excellent biomimetic of the extracellular matrix and have found great use in tissue engineering. Nanoporous monolithic hydrogels have limited mass transport, restricting diffusion of key biomolecules. Structured microbead-hydrogels overcome some of these limitations, but suffer from lack of controlled anisotropy. Here we introduce a novel method for producing architected hydrogels based on entanglement of microstrands. The microstrands are mouldable and form a porous structure which is stable in water. Entangled microstrands are useable as bioinks for 3D bioprinting, where they align during the extrusion process. Cells co-printed with the microstrands show excellent viability and augmented matrix deposition resulting in a modulus increase from 2.7 kPa to 780.2 kPa after 6 weeks of culture. Entangled microstands are a new class of bioinks with unprecedented advantages in terms of scalability, material versatility, mass transport, showing foremost outstanding properties as a bioink for 3D printed tissue grafts.

## Introduction

Microextrusion 3D bioprinting has recently been used to create functional analogues of many tissues including myocardium, skeletal muscle, liver, skin, bone, and cartilage, amongst others (*1*). Importantly, bioprinted constructs have been shown *in vivo* to possess key advantages compared to control samples made from bulk materials (*2*). The field has profited from a flurry of recent hardware and material advances, however existing bioinks have critical limitations. A major challenge is to control their flow properties to enable reproducible and accurate bioprinting with full biocompatibility. The typical bioink consists of a solution of hydrogel precursors whose rheological properties have been tuned so the bioink flows out of the nozzle in a continuous strand. The strands are then collected on a buildplate and stacked in a layer-by-layer fashion to create a 3D object. The challenge of extrusion bioprinting, referred to as a ‘race against instabilities’ (*3*), is to prevent flow and preserve the fidelity of the print until it can be stabilized by crosslinking.

To address the flow challenges of common bioinks, the field has resorted to a number of approaches. Increasing the polymer content of bioinks enhances printability (*4*), but the resulting densely crosslinked networked can reduce cell viability and spreading. Methods to print low content bioinks include 1) the addition of flow modifying fillers or additives (*5*) 2) printing within a sacrificial support material (*6*) 3) partially crosslinking the bioink before or during extrusion to increase yield stress amongst others (*7*), 4) templating using a rapidly crosslinkable material, often alginate (*8*) and 5) sequential crosslinking of individual layers directly after deposition (*9*). In general, stability of structures during printing is challenging given the high water content of most bioinks and tendency of heavy and tall structures to sag under their weight. To make matters more complex, many approaches which improve printability, have a detrimental effect on the biological properties of the material. Therefore, there is a pressing need for advanced bioinks which combine both excellent printing and bioactive/biological properties.

Microgel bioinks have been proposed as a promising strategy which could fulfill both requirements (*10, 11*). To address the flow problem, microgel bioinks have the advantage that they are not liquids and have no intrinsic flow behavior. Such bioinks also have excellent shear thinning and shear recovery properties and reasonable printing properties. They can be ‘jammed’ into a close-pack state, and held together by weak particle-particle interactions which are disrupted during extrusion in the nozzle and reformed during the post-print phase (*10*). Simultaneously, due to the inherent void space between the microgels in a close-packed state, cells within these materials have enhanced viability, spreading and migration compared to bulk monolithic materials (*12, 13*).

Current approaches to prepare and print spherical microgels for extrusion bioprinting and other biomedical applications include spraying, microfluidic, emulsion and stereolithography (*14*). All of these approaches have advantages and disadvantages, including poor scalability, the need for oils and additives, and restriction to the use of low viscous polymer solutions. Perhaps the greatest limitations of these approaches however are related to the sphericity of the microgels themselves as the close-packed lattice 1) limits interaction between individual spheres and 2) precludes any anisotropy of the structure. As a consequence, current microgel materials which are printed need to be secondarily crosslinked for stability in aqueous media and they are less suitable for anisotropic tissues like muscle and tendon where guidance of cell and matrix alignment is a critical factor.

The use of anisotropic microgels has long been explored in tissue engineering. In fact, cell-laden hydrogel microfibers have been used as an excellent tissue building block (*15,16*). Also the ability of elongated microgels (*17*) has been shown to be useful in guidance of neurons (*18*). Recently micro-ribbons produced by wet spinning and further processed into smaller parts showed the possible advantages of anisotropic gels in cartilage engineering (*19*). However a facile and cell friendly method to produce large volumes of high-aspect ratio microstrands is needed. This paper presents a new class of microgels, termed entangled microstrands, which overcomes many of the disadvantages of spherical microgel materials and demonstrates the first use of elongated microgels in 3D bioprinting.

Entangled microstrands were prepared by pressing a bulk hydrogel through a grid with micron sized-apertures to deconstruct the hydrogel into individual microstrands (Fig. 1A). The production is fast, requires no specialized equipment and can be used with wide range of pre-swollen hydrogels of arbitrary polymer content, composition, crosslinking chemistry and stiffness. Due to the simplicity of the production method, it can be upscaled to liter volumes. The void space between strands form an interconnected porous network and entangled microstrands exhibit long term stability in aqueous solutions, even without secondary crosslinking. The material is mouldable and exhibits all relevant rheological properties for 3D bioprinting. Furthermore, microstrands align in the direction of printing during extrusion through a nozzle. High fidelity 3D printing could be achieved with a wide range of materials and with polymer contents which typically require the use of support baths or additives to be printable. A bioink can be created from this material by either embedding cells *inside* the gel phase of the microstrand or depositing cells in the void space *outside* the microstrands (Fig. 1B), with both approaches providing excellent viability (>90%). Finally, we demonstrate the power of this approach for tissue engineering by 3D bioprinting of chondrocytes, showing the *in vitro* maturation of these constructs into a tissue with mechanical properties approaching that of native cartilage.

**Fig. 1.**
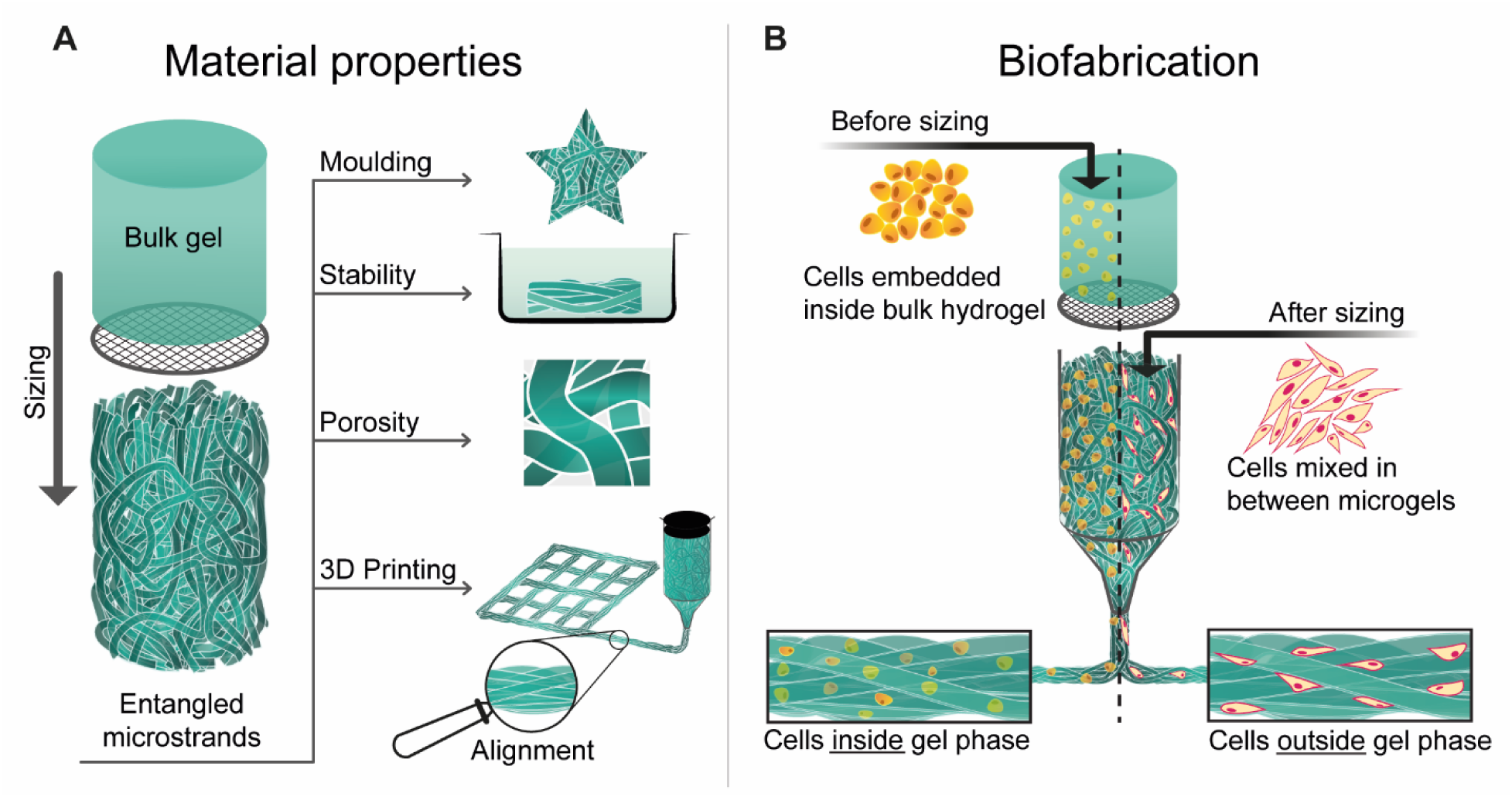
Graphical abstract: (**A**) Bulk hydrogel is mechanically extruded through a grid to size it into microstrands. In this process, microstrands randomly entangle within each other and form *entangled microstrands* a stable material with several properties, relevant for tissue engineering: mouldability, stability in aqueous solutions, porosity, printability and alignment of microstrands by extrusion. (**B**) A bioink can be prepared by embedding cells in bulk hydrogel before the sizing that results in a spatial deposition of cells inside the gel phase. Alternatively cells can be mixed in between already prepared entangled microstrands so cells occupy the space outside the gel phase.

## Results

### Entangled Microstrand Materials are Mouldable, Stable in Water and Macroporous

Here we report on a robust and versatile method for preparing ‘entangled’ microstrands. Bulk hyaluronan-methacrylate (HA-MA) hydrogels were mechanically pressed through a sieve with pores ranging from 20 to 150 microns. This process sizes the gel into microstrands, which randomly entangle within each other and make up a hydrogel material consisting of pure, high aspect ratio hydrogels. When a 2% bulk HA-MA (Degree of substitution 0.28, UV-A exposure; fig. S1) was passed through a 40 micron sieve, this resulted in a macroporous material which was visibly opaque and permeable to dyes (Fig. 2A and B, movie S1). When entangled microstrands are manipulated, single microstrands become visible (Fig. 2C). Entangled microstrands are also deformable and mouldable (Fig. 2D). Secondary crosslinking of entangled microstrands (UV-A exposure) created a macroporous structure which could be manipulated with forceps (Fig. 2E and F).

**Fig. 2.**
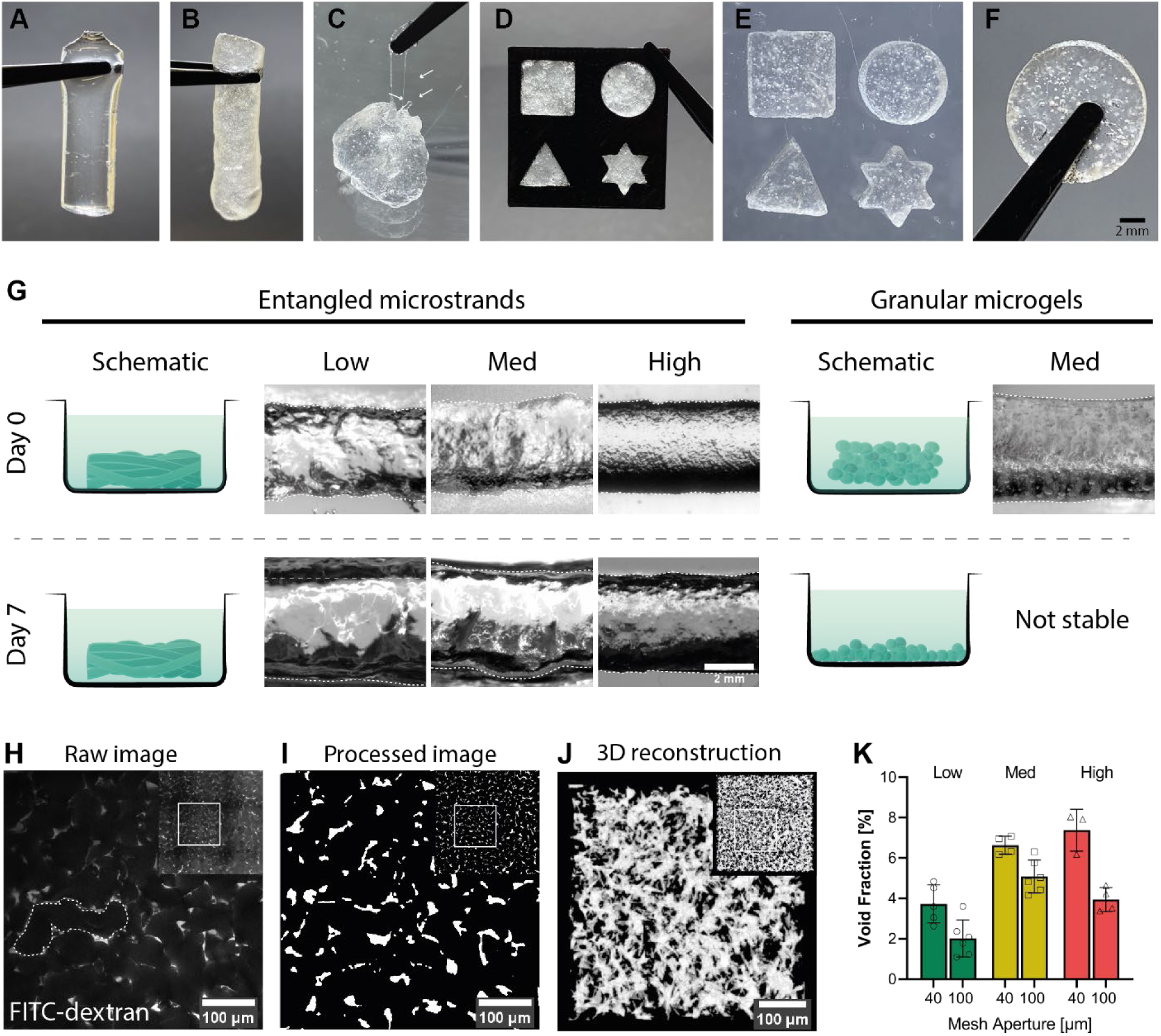
Stability and macroporosity of entangled microstrands: (**A**) crosslinked bulk HA-MA hydrogel (**B**) entangled microstrands prepared from such bulk hydrogel (**C**) when entangled microstrands are extended, single microstrands become visible (arrows) (**D**) entangled microstrands are mouldable (**E**) secondarily crosslinked custom shapes prepared by casting (**F**) secondary crosslinking tightly anneals microstrands (**G**) entangled microstrands show long-term stability in aqueous solution even without secondary crosslinking, while granular microgels loose cohesion and disintegrate (**H**) multiphoton image of entangled microstrands submerged in FITC-dextran (**I**) the same image after processing with a thresholding algorithm (**J**) 3D reconstruction of the porous network (**K**) Void fraction significantly varies with crosslinking density (F (_2, 22_) = 41.05; P<0.001) as well as mesh size (F (_1, 22_) = 48.40; P<0.001)

To investigate the universality of entangled microstrands, a range of different HA-MA bulk gels were prepared. Crosslinking of HA-MA gels was terminated at three different time points, to create bulk hydrogels with a low (*Low)*, medium (*Med*) and high (*High*) degree of crosslinking (fig. S3A). These hydrogels had significantly different mechanical properties: storage moduli (F (_2,24_) = 12164, P<0.001), compression moduli (F (_2,12_) = 34, P<0.001), maximum elongation until rupture (F (_2,6_) = 44.2, P<0.001), and swelling behavior (F (_2,14_) = 463.2, P<0.001; tab. S1 and fig S2, B to E). These three distinctively different bulk hydrogels were then further processed by sizing them through nylon meshes of two different apertures (40 and 100 microns) to create a total of 6 different variants of entangled microstrands.

Entangled microstrands had enhanced stability in aqueous medium compared to granular microgels. To compare stability, entangled microstrands and granular microgels were extruded and submerged in PBS for up to 7 days at 37°C with constant agitation (Fig. 2G). All six entangled microstrand materials (40 and 100 microns, *Low*, *Med*, *High* crosslinking) were stable for the entire period without secondary crosslinking, while all granular microgel materials based on the same hydrogel material dissociated within 1 hour of incubation (n = 3).

Porosity is a critical property of materials employed in tissue engineering as this strongly influences transport of nutrients, gas exchange and cell viability. Pore size is also relevant for blood vessel formation as well as cell migration (*20*). To assess the macroporosity of entangled microstrands, freshly prepared entangled microstrands were submerged in a fluorescent high molecular dextran dye. Fig. 2H shows a multiphoton image of the dye distribution taken within the central region of the structure, where the void space and hydrogel strand could be clearly distinguished as dextran does not penetrate inside the dense polymer network of the microstrand. Since the dye is detectable within the central region and diffusion through the hydrogel material itself is not possible, this proves the interconnectivity of the pores in entangled microstrands. To calculate the void fraction of entangled microstrands, images were processed with an algorithm (Fig. 2I) to achieve distinct transitions between microstrands and pores. A 3D reconstruction of the interconnected network can be seen in Fig. 2J while a quantification of the void fraction is displayed in Fig. 2K. Void fraction significantly varied with crosslinking density (F (_2,22_) = 41.05; P<0.001) as well as mesh size (F (_1,22_) = 48.40; P<0.001).

### Entangled Microstrand have Shear Thinning/Recovery Properties and Clear Yield Point

Entangled microstrands exhibit all relevant rheological properties necessary for extrusion 3D (bio)printing. All prepared variants of entangled microstrands showed shear thinning behavior (Fig. 3, A and B). To identify the yield stress necessary to induce flow different methods can be employed (*21, 22*). In this study we used the crossover of G’ and G’’ to identify the flow point. Required stress to reach crossover was lowest in High samples and highest in *Low* samples for both mesh apertures (40 µm: *Low* = 450 Pa, *Med* = 409 Pa, High = 141 Pa; 100 µm: *Low* = 659 Pa, *Med* = 559 Pa, *High* = 225 Pa). To simulate the printing process, shear recovery tests based on oscillatory strain sweeps with cycles of high and low strain were conducted. At low strains, microstrands exhibited a solid-like elastic behavior (G’ > G’’) that rapidly changed into a liquid-like viscous behavior (G’ < G’’) when high strains were applied. These transitions are crucial for high quality 3D printing as it ensures even material flow during extrusion and shape retention upon deposition on the collector plate.

**Fig. 3.**
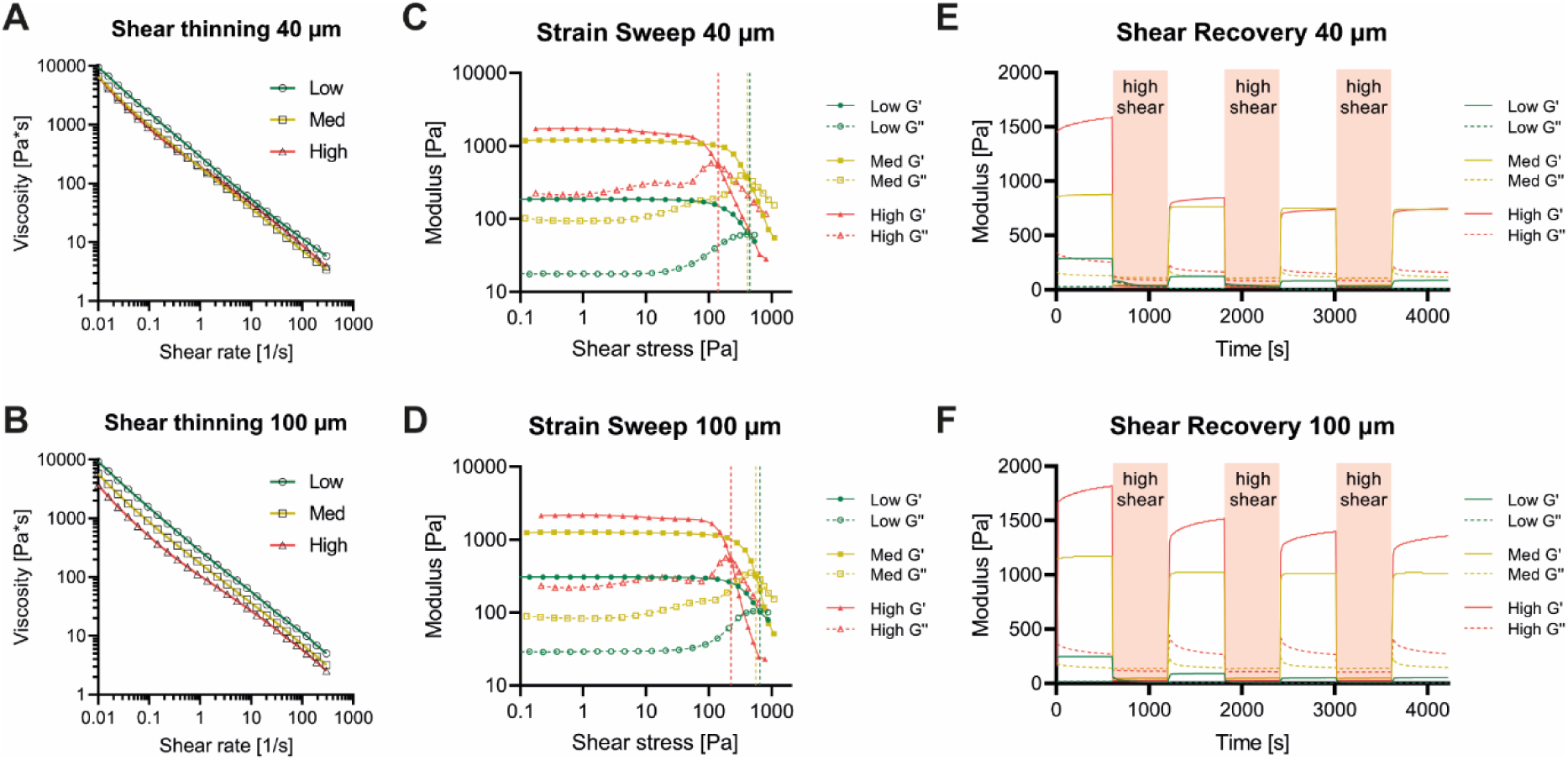
Rheological characterization of HA-MA entangled microstrands: (**A - B**) entangled microstrands created by sizing with a grid with aperture size of 40 and 100 µm exhibit shear thinning behavior (**C - D**) clear flow points can be determined (Crossover points for 40 µm samples: *Low* = 450 Pa, *Med* = 409 Pa, *High* = 141 Pa; 100 µm samples *Low* = 659 Pa, *Med* = 559 Pa, *High* = 225 Pa) (**E - F**) when subjected to repeated cycles of low and high shear, shear thinning and shear recovery behavior can be observed for all conditions

### Entangled Microstands have Excellent Printability and Anisotropic Structure When Printed

To test the printability of entangled microstrands, a two-layered grid structure was printed (Fig. 4A). All six variants were printed with good shape retention and printing resolution. Anomalies arose for lower crosslinked samples when sharp edges were printed. This phenomenon was especially pronounced in *Low* crosslinked samples (40 µm) where sharp edges have a very rounded appearance and the filament was dragged away during printing. This problem can be explained with the stronger mechanical strength of the microstrands compared to a typical polymer solution. The printed filament extends from the printing nozzle to the deposited filament of the printed structure. In a typical polymer solution, the filament collapses as soon as the nozzle moves away from the printed construct. In *Low* crosslinked microstrand samples, however, microstrands spanned from the printed construct up into the printing nozzle and therefore the printing filament did not rupture but was rather dragged to the new printing position. Since more crosslinked samples like *Med* and *High* were more brittle and ruptured at lower elongation distances, the printing accuracy was better in these samples.

**Fig. 4.**
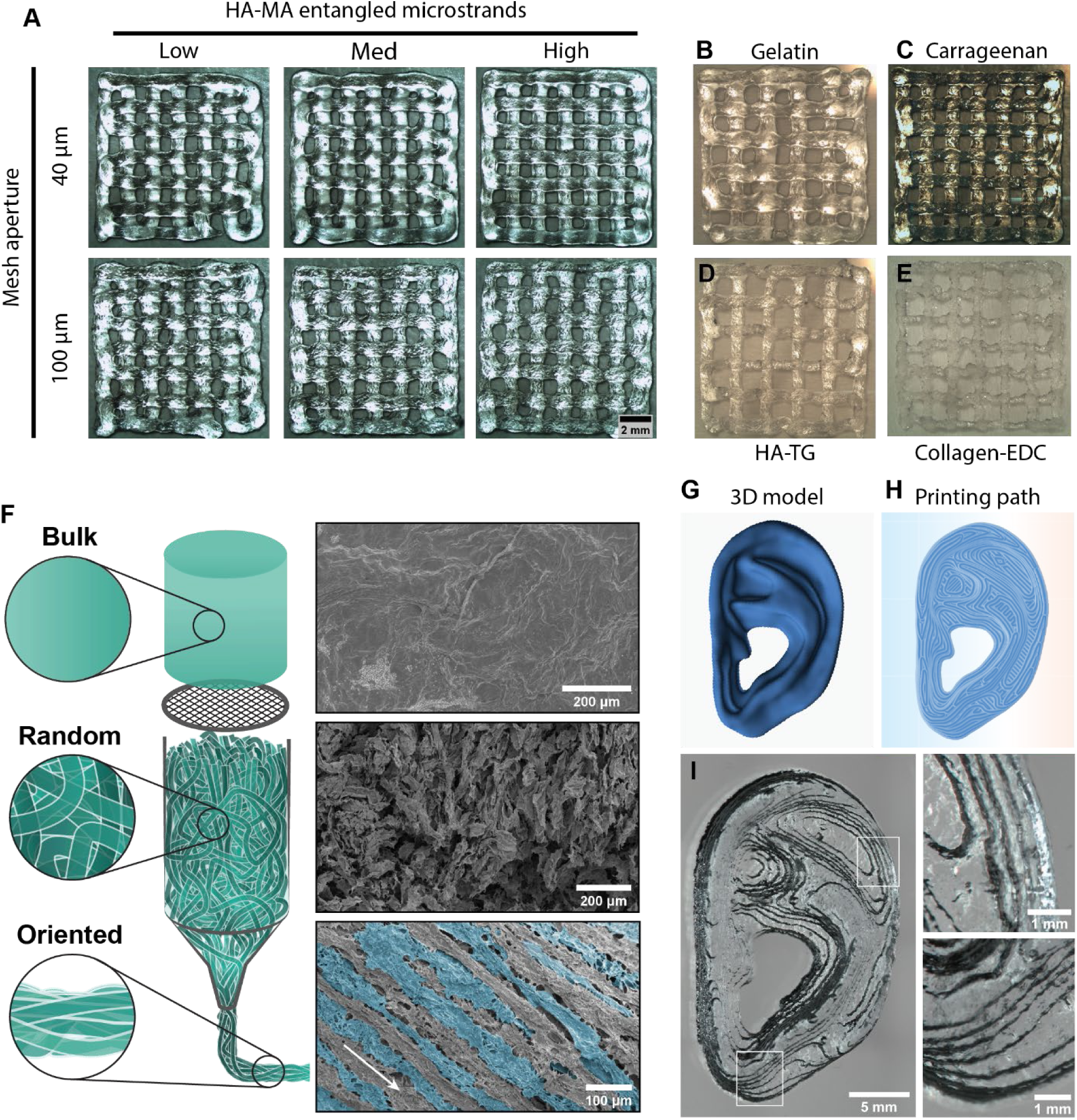
Entangled microstrands are printable and align during extrusion: (**A**) Different HA-MA entangled microstrands printed in a grid structure (**B**) Entangled microstrands prepared and printed from i-carrageenan (ionic crosslinking), (**C**) gelatin (thermal crosslinking), (**D**) HA-TG (enzymatic crosslinking), (**E**) Collagen-EDC (carbodiimide crosslinking) (**F**) SEM images of bulk gel, freshly prepared entangled microstrands in random orientation and aligned microstrands after extrusion through a printing nozzle. The arrow indicates the direction of extrusion (**G**) 3D model and (**H**) printing path of a human shaped ear (**I**) 3D printed with entangled microstrands prepared from carrageenan

To demonstrate the universality of this approach, microstrands were created from five common biomaterials. Gelatin, a thermogelling denatured form of collagen, is widely used in the tissue engineering field in its unmodified form or as a photo-activated material called gelatin-methacrylol (gelMA) (*23*). ι-carrageenan is a highly sulfated polysaccharide that forms ionic crosslinks upon the addition of monovalent as well as divalent cations (*24*). Hyaluronan hydrogels were either crosslinked enzymatically by activated factor XIII (*25*) or chemically by divinyl sulfone (*26*). Finally, collagen, a material prevalent in tissue engineering because of its abundance in the extracellular matrix of many tissues was crosslinked with carbodiimide chemistry (*27*). Although derived from a wide range of materials and crosslinking methods, all bulk hydrogels were successfully sized and the entangled microstrands could be 3D printed in a grid shape following the same protocol as used for the HA-MA microstrands (Fig. 4, B to E, Fig S3, A and B).

To demonstrate the power of 3D printing of these bioprinting materials, a biologically relevant structure (ear, Fig. 4G) was printed with entangled microstrands prepared from bulk carrageenan. The printing fidelity was such that the individual layers were strongly visible, clearly mirroring the printing path (Fig. 4, H and I).

Anisotropy in tissues like muscle, tendon or nerves is difficult to replicate with conventional tissue engineering approaches. Several studies have shown the possibility to align (nano-)fibers with the direction of flow (*28*). Applied to the field of 3D bioprinting, this allows the creation of materials with aligned fibers embedded within the hydrogel. It is unclear however, to which degree cells can interact and align according to these fibers embedded within a bulk hydrogel.

To investigate the ability of entangled microstrands to align during 3D printing, HA-MA microstrands were extruded through a 410 µm conical printing nozzle and their structure observed with scanning electron microscopy (Fig 4F). Clear differences between bulk gel, freshly prepared microstrands and 3D printed microstrands were apparent. Hydrogel had a smooth surface, whereas freshly prepared microstrands were randomly entangled within each other and had no clear orientation. Extrusion through a printing nozzle, however, aligned the microstrands in the direction of printing. A deposited filament therefore consisted of a multitude of individual microstrands aligned in the print direction.

### Cellular Entangled Microstrands Lead to Rapid Tissue Maturation

To explore the potential for 3D bioprinting with entangled microstrands, two possible cell delivery approaches were investigated (Fig 1B). Firstly, cells were embedded inside the bulk hydrogel before the hydrogel was sized into microstrands to create cell-laden microstrands (*Inside*). Alternatively, entangled microstrands were mixed with a cell suspension, leaving cells to occupy the void space between microstrands (*Outside*).

To investigate the *Inside* approach, C2C12 cells were embedded within a solution of 2% gelatin and 2% GelMA und subsequently sized (Cells inside gel phase, Fig 5A). A significant, but minor drop of viability was observed for cells embedded in bulk hydrogel compared to freshly trypsinized cells (2D 98.2±0.6%; Bulk = 92.6±1.6%; t(4)=4.6; P < 0.01). The viability of cells in hydrogels that have been sized into entangled microstrands was high (40 µm = 93.2±0.8%, 100 µm = 94.1±0.7%) and no significant difference in viability was found between either of the sized samples or the bulk gel controls (F (_2,6_) = 0.9353, P=0.443).

**Fig. 5.**
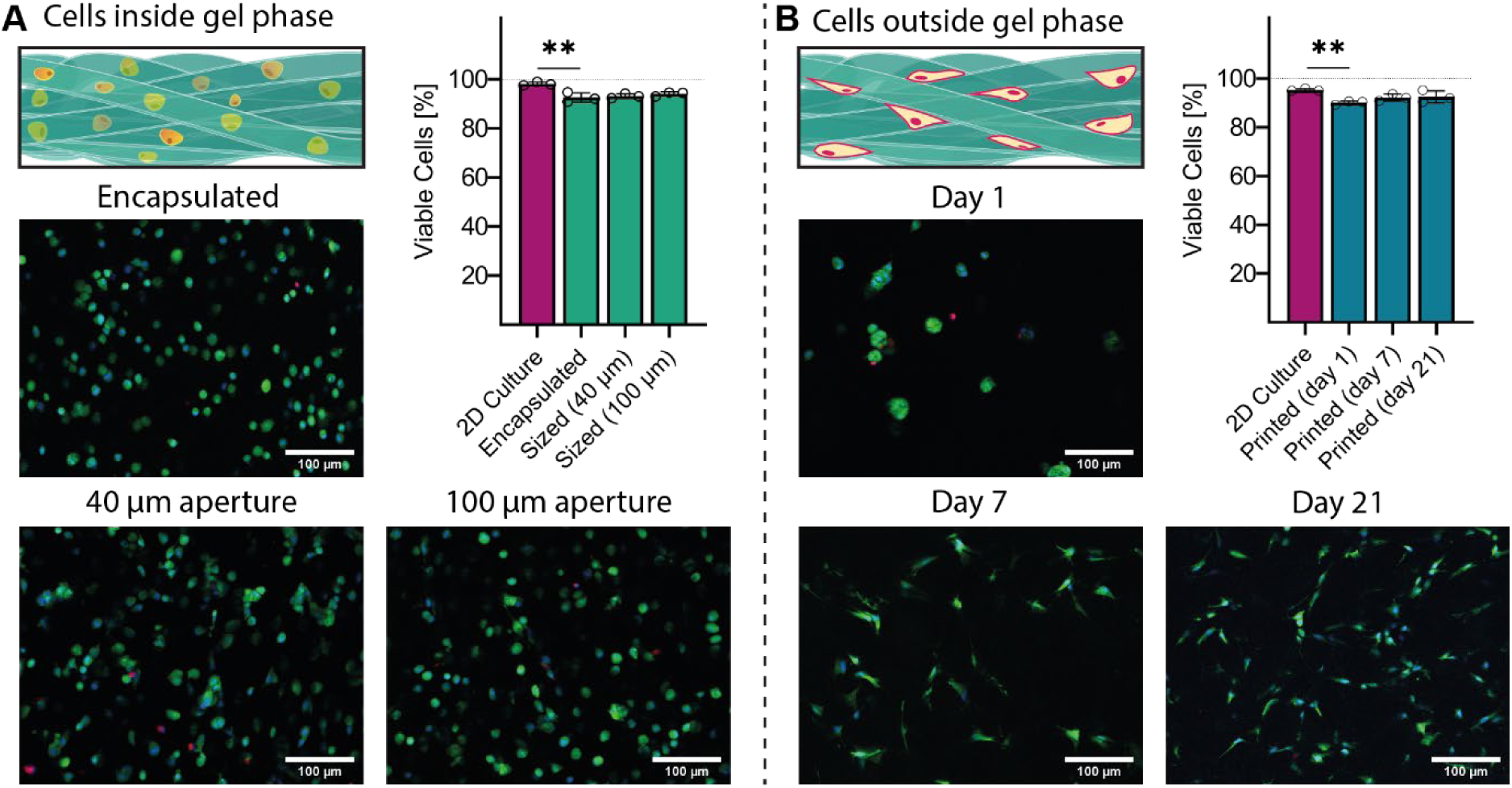
Entangled microstrands allow 3D bioprinting with high cell viability: (**A**) Cells can be embedded in bulk hydrogel and subsequently sized into cell-laden microstrands without negative impact on viability. Fluorescent images of cell-laden entangled microstrands bioink (green = alive, red = dead) show a healthy population of cells and quantification reveal a significant, but minor drop on viability when cells are encapsulated but not when sized through the grid (**B**) Alternatively, cells can be mixed with already prepared entangled microstrands and 3D bioprinted with minimal impact on cell viability. Fluorescent images show high viability of cells after the printing process and cells adhering to the outer surface of entangled microstrands, proliferating and occupying the porous space at Day 7 and Day 21.

To test the effect of pore space of entangled microstrands on matrix production, bovine chondrocytes were combined with HA-MA microstrands and 3D bioprinted into cylindrical discs (Fig. 5B). In this approach, chondrocytes were within the void fraction of the entangled microstrands and not embedded within the polymer network of the hydrogel itself (*outside*). In a fluorescent Live/Dead staining, chondrocytes showed a rounded phenotype after 3D bioprinting, but displayed a spread phenotype after 7 and 21 days in culture (Fig. 5B). Cell viability was above 90% for the entire duration of the experiment (Pre-printing = 95.3±0.5%; Day 1 = 90.1±0.6%; Day 7 = 92.3±1.1%, Day 21 = 92.6±2%). Even though there was a significant drop in cell viability when bioprinted cells are compared to the original cell population (t(4)=9.6; P < 0.001), cell viability decreased by just 5.2±1.2% (Pre-Printing to Day 0). Cell viability continued to stay high and no statistical difference in viability was found between printed cells at day 1 and 7 or 21 (F(_2,6_) = 1.873, P=0.233). When compared to literature, the percentage of viable cells is extremely high for a microextrusion based approach and superior when compared to most biofabrication methods (*29*). This is also true when directly compared to alternative microgel techniques in which cells are encapsulated within hydrogel microbeads. Although cells are protected from shear stress in such a setup and no drop in viability is observed after 3D bioprinting, microfluidic preparation impairs viability at similar or greater levels and usually results in reported viabilities of 70-90% (*10, 11*).

With increasing proliferation at later time points, the spatial distribution of cells became more apparent. Cells grew and occupied the void space between microstrands and therefore the pattern of the entangled microstrands and the porous network within became visible (Fig. 5B).

To investigate the use of entangled microstrands as a platform for cartilage tissue engineering, bioprinted discs were cultured *in vitro* for up to 6 weeks. Immediately after fabrication, bioprinted scaffolds were transparent and slightly cloudy (Fig. 6B). Turbidity in these scaffolds can be attributed to the presence of cells, since acellular printed samples were transparent. After 6 week culture, appearance changed to a cartilage-like opaque white, indicating deposition of a dense extracellular matrix (Fig. 6C). Cultured tissue constructs were fixed and histologically stained for cartilage specific markers. A representative sample of the population (n = 6) is depicted in Fig. 6A. Staining of Safranin O showed an increased intensity with time, indicating strong proteoglycan content in the samples. Staining with Hematoxylin & Eosin (H&E) resulted in a contrast of the interface between stained cells and unstained entangled microstrands. While high levels of collagen type I were detected 3 weeks after fabrication, collagen I staining was reduced after 6 weeks of maturation. Collagen type II was present after 3 weeks, but restricted to the void space in between microstrands. After 6 weeks of culture, collagen II staining intensified and showed deposition inside the void space, as well as the hydrogel network of the entangled microstrands. The difference in staining between time-points was particularly striking in the outer ~400 µm of the sample (Fig 6A). After 3 weeks, stainings for cartilage-ECM markers as well as abundance of cells in the outer area of the sample was lower compared to the central part of the scaffold. This phenomenon inverted after 6 weeks of culture as there was a very dense deposition of ECM as well as cells in the outermost part of the scaffold. We hypothesize that this occurrence might be linked to the degradation of the hydrogel microstrands. At the 6 weeks time-point, cells in the outermost part did not show the initial pattern of thin lines and small clusters also found in Live/Dead stainings anymore, but had a more homogeneous distribution. This indicates that cells were able to migrate into the space previously occupied by the hydrogel microstrands.

**Fig. 6.**
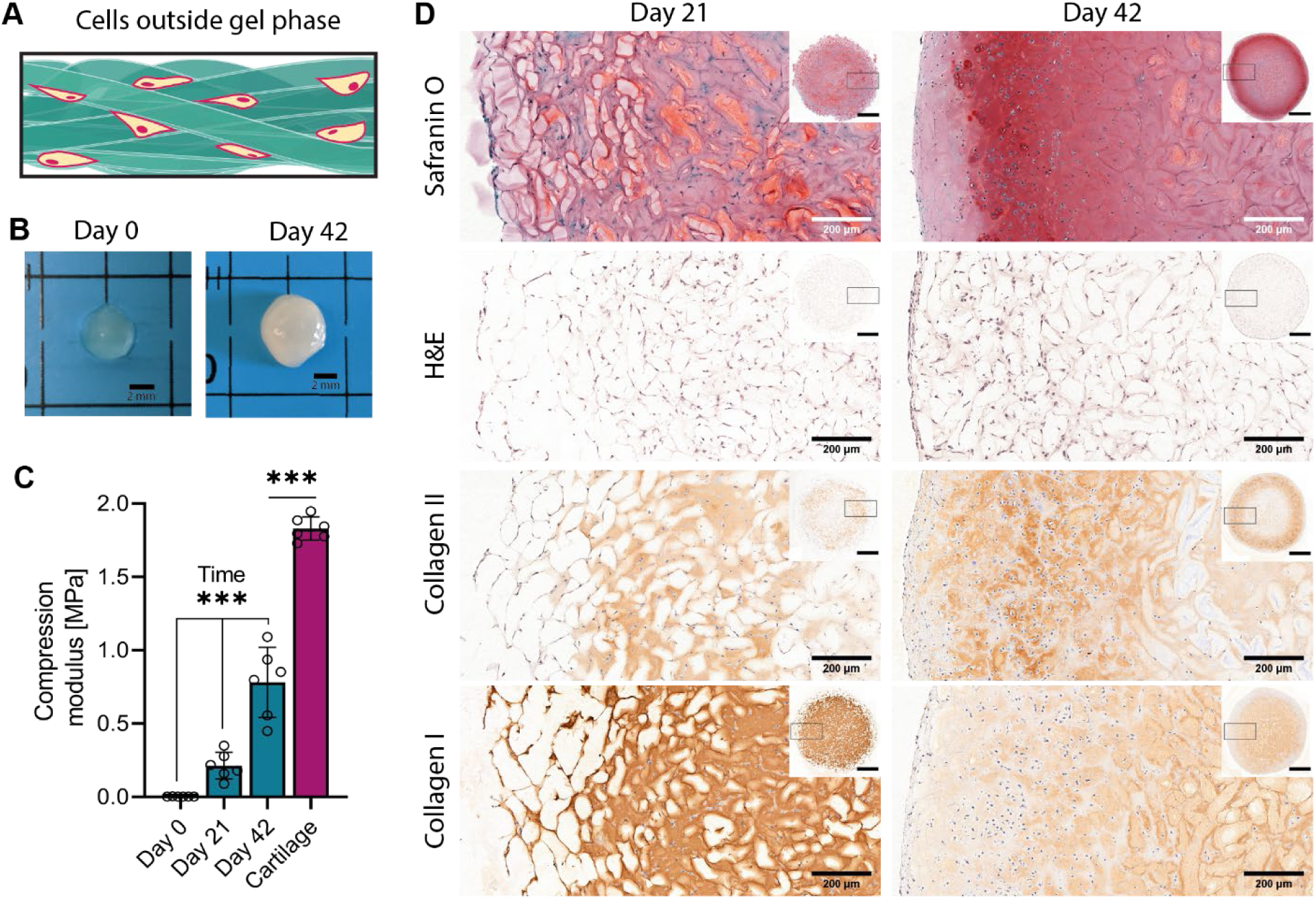
Entangled microstrands mature into cartilage-like tissue: (**A**) In this experiment, bovine chondrocytes were mixed with already prepared HA-MA microstrands (**B**) appearance of 3D bioprinted discs changed from transparent (day 0) to shiny-white (day 42) (**C**) comparison of compression modulus of freshly bioprinted entangled microstrands, after *in vitro* culture and healthy articular cartilage (**D**) histological staining to highlight the strong deposition of cartilaginous matrix entangled microstrand scaffold directly after printing and after 6 weeks of culture

The mechanical properties of cartilage tissue is highly important for its functions. Especially the ability to sustain load is one of the most crucial ones. Compression modulus of freshly bioprinted samples was very low (2.7±0.3 kPa), but showed a significant increase over time (F* (_2,15_) = 44.38; P < 0.001), reaching 212±83.7 kPa after 3 weeks and 780.2±218.4 kPa after 6 weeks. While this was still significantly lower when compared to native articular bovine cartilage which has a compression modulus of 1829.8±72 kPa (T(10)=10.2; P < 0.001), this is a remarkable increase. Since the material used in this study had very little resistance to compression on its own, change in compression modulus could exclusively attributed to the abundant deposition of extracellular matrix.

## Discussion

Prior studies and recent reviews have documented the strength and potential of microgels for tissue engineering (*2, 13*). While this method allows for great versatility, the spherical nature of microgels used in those approaches have certain limitations. Due to the inherent surface to volume ratio of spheres, interaction is low, which can lead to poor long-term stability of constructs. For the same reason, aligned structures are highly challenging to prepare with spherical microgels, as this would require manipulation of the arrangements of microgels at a level, which has not been demonstrated yet.

With our simple but effective approach to produce entangled microstrands, it is possible to mould or print stable, porous hydrogel structures. The microstrands that make up this hydrogel can be aligned by extrusion through a nozzle, giving orientation to printed ‘architected’ constructs. We demonstrated printability of mechanically different HA-MA gels as well as a wide range of other hydrogels spanning different physical and chemical crosslinked hydrogels. This indicates that entangled microstrands are applicable to a wide range of possible hydrogels. These hydrogels are also clinically-relevant as hyaluronan is widely used as injectable dermal fillers and as viscosupplements for joint pain. Some of these medical products are crosslinked with DVS, similar to the way it was done in this study (*26*).

We showed successful 3D bioprinting when entangled microstrands prepared from HA-MA were combined with bovine chondrocytes and printed with excellent viability.

Furthermore, chondrocytes within the 3D printed constructs deposited copious cartilaginous extracellular matrix which translated into a significant increase in stiffness.

Void fraction of the macroporous network provided by entangled microstrands is low compared to those provided by spherical microgels. However, our data shows that the porous network is interconnected and that cells can utilize the porous network for proliferation and deposition of extracellular matrix. Since some tissues might require higher void-fraction, grids used in this study could be substituted by photo-etched metal plates with apertures of custom cross section e.g. star shaped, to further increase distance between microgels.

Going one step further, multimaterial approaches are the next step to introduce heterogeneity in microgel systems (*2, 13*). For entangled microstrands this is relevant at two different levels. On the one side, the simple production of entangled microstrands allows combinations of hydrogel precursors to be prepared as one bulk gel and sized into entangled microstrands consisting of multiple materials. Following this approach, double network hydrogel microstrands can be prepared or microstrands in which a sacrificial gel (e.g. gelatin) gives printability to any other polymer precursor.

In a complimentary approach, microstrands produced from different bulk gels can be combined. Even though a homogeneous mixture of different microstrands without any damage to the material might be challenging to prepare, multibarrel syringes with intricate outlets might solve this problem. Application of such a complex microstrands system would be manifold, ranging from directed gradients of microstrands loaded with biologicals to defined control over porosity via sacrificial microstrands.

In conclusion, we presented a new class of bioinks, called entangled microstrands, which unify many crucial properties needed for tissue engineering scaffolds. We successfully demonstrated the use of this material to facilitate rapid tissue maturation in the presence of cells. Preparation and processing of entangled microstrands is facile and transferable to a wide range of hydrogel systems, including clinically relevant hydrogels. These advantages together suggest that the translation of these materials for medical applications will be more straightforward compared to other commonly used bioinks and bioprinting materials.

## Materials and Methods

All chemicals were purchased from Sigma-Aldrich unless stated otherwise.

### Polymer Synthesis and gel preparation

#### HA-MA

Hyaluronic acid (1 gram, HTL Biotechnology) was dissolved in 400 ml ultrapure water and kept at 4°C overnight to ensure complete dissolution. Ice-cold DMF (267 ml) was added under continuous stirring. To start the reaction methacrylic anhydride (2370 µl) was added and the pH kept between 8 - 9 through the addition of 10 M NaOH for 4 hours. Solid sodium chloride was dissolved in the solution to achieve a concentration of 0.5 M and the polymer was subsequently precipitated with ethanol (Merck). The precipitate was washed with ethanol, dried and dissolved in ultrapure water. Solution was purified by diafiltration (Äkta 3, 10 NMWC hollow fiber). The purified product, hyaluronic acid methacrylate (HA-MA) was lyophilized dissolved in deuterium (Cambridge Isotope Laboratories) and characterized by ^1^H NMR spectroscopy and stored at −20°C in the dark until used.

NMR spectra were recorded at room temperature on a Bruker AV-NEO 600 MHz spectrometer equipped with a TCI cryo probe. Spectra were obtained with 1024 scans using a 5 s recycle delay. To determine the degree of substitution, we compared the ratio of the sum of the integrated peaks of the methacrylate protons (peaks at ~6.1 and ~5.6) and the integrated peak of the methyl protons of HA (~1.9 ppm).

For gel preparation, HA-MA was dispersed in PBS and kept at 4°C until complete dissolution. HA-MA solution was mixed with a 1% lithium phenyl-2,4,6-trimethylbenzoylphosphinate (LAP) stock solution to create a 2% HA-MA and 0.05% LAP solution and crosslinked by controlled photoexposure in the UV-A range (Omnicure Series 100, 400 nm wavelength, 9.55 mW cm^−2^).

#### Gelatin Methacrylol (GelMA)

Gelatin type A was dissolved in PBS at pH 7.4 and warmed up to 50°C under vigorous stirring. Total used MA volume was split into five and after every addition, pH was adjusted with NaOH and the solution left to react for 30 minutes. After the last addition, reaction was diluted 2 fold and left to react for another 30 minutes. Product was cleaned by subsequent dialysis (10-12 kDa Cutoff) against ultrapure water for 4 days. Solution was filtered, lyophilized and stored at −20°C until use.

For gel preparation, GelMA was dissolved in 70°C hot PBS. GelMA solution was mixed with LAP stock solution (1%) to achieve a final concentration of 2% GelMA and 0.1% LAP. Solution was photocrosslinked by controlled photoexposure in the UV-A range.

#### Hyaluronic acid transglutaminase (HA-TG)

For HA-TG hydrogel precursors, two different batches of HA were substituted with reactive glutamine (HA-TG/Gln) and lysine (HA-TG/Lys) residues respectively following published protocols (*25*).

For gel preparation, HA-TG/Lys and HA-TG/Gln were dissolved in TBS buffer (150 mM NaCl, 40 mM CaCl2, 50 mM TRIS, pH 7.6) and combined at equal volume to form HA-TG solution. To initiate gelation, a solution of thrombin (Baxter, 500 U / ml) and factor XIII (Fibrogammin, CSL Behring, 200 U / ml) was added to form a gel with final concentrations of 3% HA-TG.

#### ι-carrageenan (ι-CRG)

300 mg of *ι*-carrageenan particles (Genuvisco CG-131, GP Kelco) were added to 10 ml of 4°C cold buffer solution (150 mM KCl, 20 mM HEPES, pH 7.4) to allow hydration of particles. Dispersion was then heated to 80°C, stirred until complete dissolution, and transferred into a 10 ml syringe. Solution was cooled down and stored at 4°C to form a 3% (w/v) gel.

#### Gelatin

300 mg of gelatin particles from porcine skin (type A) were added to 10 ml of 4°C cold PBS to hydration with subsequent heating to 70°C until complete dissolution. The solution was transferred into a 10 ml syringe and cooled down to 4°C to form a 3% (w/v) bulk gel. To ensure reproducible results, gelatin solution were stored at 4°C for 24 h to minimize variances due to the hardening of gelatin gels.

#### Hyaluronic acid divinyl sulfone (HA-DVS)

A solution of 3% (w/v) hyaluronic acid, 3% (w/v) NaCl and 0.2 M NaOH was prepared and stirred vigorously until complete dissolution of hyaluronic acid. Divinylsulfone was added to the solution to equal the amounts of hyaluronic acid (w/w). Solution was mixed to ensure a homogeneous distribution and left to gel for 3 hours at room temperature. Gel was then washed for 2 days in deionized water and used for experiments.

#### Carbodiimide crosslinked collagen (Collagen-EDC)

Two different concentrations of Type I collagen solution (5 mg / ml, Symatese; 80 mg / ml, 3dbio) were mixed on ice to achieve a final concentration of 20 mg / ml. An equal amount (w/w) of 3,3’-Dithiobis(propionohydrazide) (DTPHY) was directly dissolved in this solution and 6 times excess of 1-Ethyl-3-(3-dimethylaminopropyl)carbodiimide (EDC) was first dissolved in MES buffer and subsequently added to the solution. Solution was mixed and left to react at 4°C over night to form a stable gel.

### Preparation of entangled microstrands (sizing)

To prepare entangled microstrands, bulk hydrogels were prepared inside a 10 ml syringe and manually pressed through a nylon sieve (Millipore, Filter code: NY41 for 40 µm, NY1H for 100 µm) and directly used for experiments.

### Rheology

To assess rheological properties of the samples, all measurements were conducted on an Anton Paar MCT 301 rheometer equipped with a 20 mm parallel plate geometry at 25°C in a humid atmosphere with a gap distance of 1 mm. Rheological properties of samples were examined by oscillatory shear sweeps (1% strain, 1-100 Hz), ramped shear rate (0.01 - 50 1/s) and strain sweeps (1 Hz, 0.01-1% strain) to evaluate storage and loss modulus, yield point and shear thinning behavior. To investigate shear recovery properties, samples were exposed to repeating cycles of alternating phases of strain (1 Hz, 1% and 500% strain).

### Compression modulus measurements

Disc shaped test specimen with 8 mm diameter and 2 mm height were stamped out of the bulk gel for acellular samples. For cellular samples, bioprinted constructs were used and exact dimension measured before testing. Samples were tested by unconfined compression using a texture analyzer (TA.XTplus, Stable Microsystems). A 500 g load cell and a flat plate probe with a diameter of 15 mm were used. Samples were compressed to a final strain of 15% at a rate of 0.01 mm/s. The compression modulus was calculated from the slope of the linear first 3% of the stress-strain curve.

### Elastic Modulus

Bulk gels with defined crosslinking were prepared and dumbbell shaped specimens according to ISO 527-2-5B were stamped out. Specimen were attached in a custom clamp system, and elongated until failure at a rate of 0.01 mm / s.

### Swelling

Bulk hydrogel discs were prepared and weighted before immersion in PBS to induce swelling. For each time point, samples were removed from the PBS, blotted with a tissue to remove excess PBS and weighted. The degree of swelling was calculated using following equation.

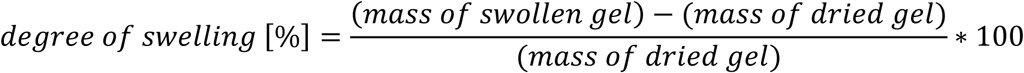

### 3D Printing

Entangled microstrands were loaded into printing cartridges (Nordson EFD) and printed through a 410 µm conical needle (Nordson EFD) with a pneumatic driven extrusion 3D bioprinter (3D Discovery, RegenHU). 3D models for grid and disc structures were created with OpenSCAD version 2015.03-2. 3D models were processed with Slic3r version 1.3.0 dev to create machine code (G-Code).

### Scanning electron microscopy (SEM)

Entangled microstrands were extruded as straight lines through a 410 µm conical needle and collected on a glass plate. Bulk HA-MA hydrogel and freshly prepared entangled microstrands were also prepared as control. All samples were frozen in liquid nitrogen and lyophilized. For SEM analyses, the lyophilized samples were coated using Pt/Pd (80/20) at a thickness of 10 nm by a sputter coater (CCU-010 HV, Safematic). The imaging was performed using a SEM instrument (JSM-7100, JEOL).

### Macroporosity

Entangled inks were prepared as described and submerged in PBS containing a high molecular weight, fluorescent FITC-dextran (average molecular weight of 500 kDa). Entangled microstrands were then imaged by two-photon microscopy (SP8, Leica).

### Stability

Entangled microstrands were prepared as described, cut into cylinders and transferred into well plates. Samples were submerged into PBS for up to 7 days and PBS was removed and exchanged after 5 min, 1 hour, 24 hours and 7 days.

### Cell laden microstrands (*inside*)

#### C2C12 mouse cell line

C2C12 mouse immortalized myoblasts were obtained from ATCC.

Cells were cultured in a humidified atmosphere with 5% CO_2_ at 37°C in Dulbecco’s modified Eagle’s medium (DMEM GlutaMAX, Gibco) with 10% fetal bovine serum (FBS, Gibco) and 10 µg/ml of gentamycin sulfate (Gibco). Cells were passaged at 90% confluence and detached by 0.25% Tripsin/EDTA (Gibco).

Encapsulation solution (2% gelatin, 2% GelMA and 0.05% LAP) was prepared and kept at 37°C to avoid solidification. Freshly detached C2C12 were added to the solution and gently mixed by continuous pipetting. After homogeneous distribution was achieved, solution was transferred into a syringe and cooled in an ice bath for 1 hour, while the syringe was constantly rotated for the first 5 minutes to avoid sedimentation. After gelation period, cell containing bulk gel was pressed through a nylon grid (sized), secondarily crosslinked by photoexposure in the UV-A range and kept in culture media.

### Bioprinting of entangled microstrands (*outside*)

#### Bovine chondrocytes

Primary articular chondrocytes were isolated from the femoral cartilage of 6 month old calves obtained from the local slaughterhouse. Cartilage from the medial and lateral condyle was harvested, minced and digested by 0.1% collagenase solution (from *Clostridium histolyticum*) over night. Cells were cultured in a humidified atmosphere with 5% CO_2_ at 37°C and high glucose Dulbecco’s modified Eagle’s medium (DMEM, Gibco) supplemented with 10% (v/v) FBS (Gibco), 50 µg / ml L-ascorbic acid 2-phosphate sesquimagnesium salt hydrate and 10 µg / ml gentamycin sulfate (Gibco). Cells were passaged at 90% confluence by detachment with 0.25% trypsin/EDTA (Gibco) and used for experiments at passage 3.

To prepare bioink, entangled microstrands and dense solution of bovine chondrocytes (100 × 10^6^ cells / ml) were loaded in separate chambers of a double barrel syringe with a chamber ratio of 10:1 (Medmix). The two components were mixed by extrusion through a static mixing element (Medmix) to prepare cell-laden entangled microstrands. Bioink was transferred into a printing cartridge (Nordson EFD) and printed into discs (d = 5 mm, h = 2 mm). To ensure long-term shape fidelity, microstrands were annealed by 15 seconds of UV-A exposure. Constructs were cultured in high glucose DMEM supplemented with 1% (v/v) ITS liquid media supplement (Fisher Scientific), 40 µg / ml proline, 50 µg / ml ascorbic acid, 10 µg / ml gentamycin sulfate and 10 ng / ml TGF-β3 (Preprotech) for up to 6 weeks with full media change three times a week.

### Live/Dead Staining

Bioprinted constructs were cut in half, washed with phenol-free DMEM (Gibco) and stained with 0.5 µg /ml propidium iodide, 0.008 mM calcein AM and 5 µg /ml Hoechst 33342 for 20 minutes and imaged with fluorescent light microscopy (ZEIS, Axio Observer Z1). Z-stack images spanning 100 µm were acquired from the center of the scaffold and analyzed with FIJI. Experiment was done in triplicates, with the viability of each sample averaged over three pictures of randomly chosen positions inside the center of the hydrogel.

### Histological evaluation

Samples for histology were fixed in paraformaldehyde for 2 hours, dehydrated and paraffinized (LogosJ, Milestone). Paraffin blocks were cut with a microtome in 5 µm thick sections, dried deparaffinized and hydrated. Tissue sections were stained with SafraninO, hematoxylin and eosin (H&E), Picosirius red and Alizarin Red according to standard protocols.

For colorimetric, immunohistochemical stainings of collagen type I and II, sections were first digested in a 1200 U/ml hyaluronidase solution (from *Streptococcus equi*) for 30 minutes at 37°C. Sections were then washed and blocked with 5% normal goat serum (NGS) in PBS for 1 hour at room temperature. Subsequently, slides were blotted and primary antibody in 1 % NGS was added and left overnight in humidified atmosphere to avoid drying. Anti-collagen type I antibody (mouse, Abcam #ab6308) was used at 1:1500 dilution, while anti-collagen type II antibody (mouse, DSHB #II-II6B3) was used at 1:200 dilution. On the following day, sections were washed with PBS and treated with 0.3% H_2_O_2_ to quench any endogenous peroxidase or pseudoperoxidase activity to prevent non-specific signals. After an additional washing step, secondary antibody (goat, anti-mouse IgG (HRP), Abcam #ab6789) in 1% NGS solution was added and left under humidified atmosphere for 1 hour. Sections were washed again and counterstained by Mayer’s hematoxylin solution. All samples were mounted and coverslipped with resinous mounting media (Eukitt) before imaging with a Pannoramic 250 histology slide scanner from 3D Histech.

### Statistical analysis

Statistical analysis was conducted with GraphPad Prism (v. 8.2.0 (425)) and statistical significance was assumed for P < 0.05. For acellular samples, storage modulus, elongation, compression and shear thinning between samples were compared with one way analysis of variance (ANOVA) with a Tukey post hoc test. For swelling of samples a mixed-effects model with Geisser Greenhouse correction was used.

The influence of mesh size and crosslinking degree on the formation of macroporosity was investigated with a two way ANOVA. To analyze the viability of encapsulated and bioprinted cells, one-way ANOVA was used. To compare viability of freshly trypsinized cells with encapsulated ones, an unpaired, two-tailed T-Test was conducted.

Mechanical properties of cellular, tissue engineered constructs were compared with a Brown-Forsythe and Welche ANOVA test. Additionally, Day 42 samples were compared to native cartilage with an unpaired, two tailed T-Test.

## Supporting information

microstrands video

## H2: Supplementary Materials

Results

Sup. 1. NMR spectra of synthesized HA-MA

Sup. 2. Mechanical properties of HA-MA gels

Sup. 3. 3D printed GelMA and HA-DVS

Table S1. Mechanical Properties of Different HA-MA Bulk Gels

Movie S1. Porosity of entangled microstrands

## General

We want to thank Dr. Alvar Diego Gossert and Philipp Rössler for their help with the NMR-spectra acquisition. We also want to thank David Fercher, Dominic Rütsche and Riccardo Rizzo for their help in the preparation and synthesis of HA-TG, Collagen-EDC and GelMA.

## Author contributions

Describe the contributions of each author (use initials) to the paper. B.K., M.L. and M.Z. designed the research, B.K. and M.Z. analyzed the data and wrote the paper, B.K., A.B., M.L., Y.T. and E.T. performed the research.

---

**Sup. 1.**
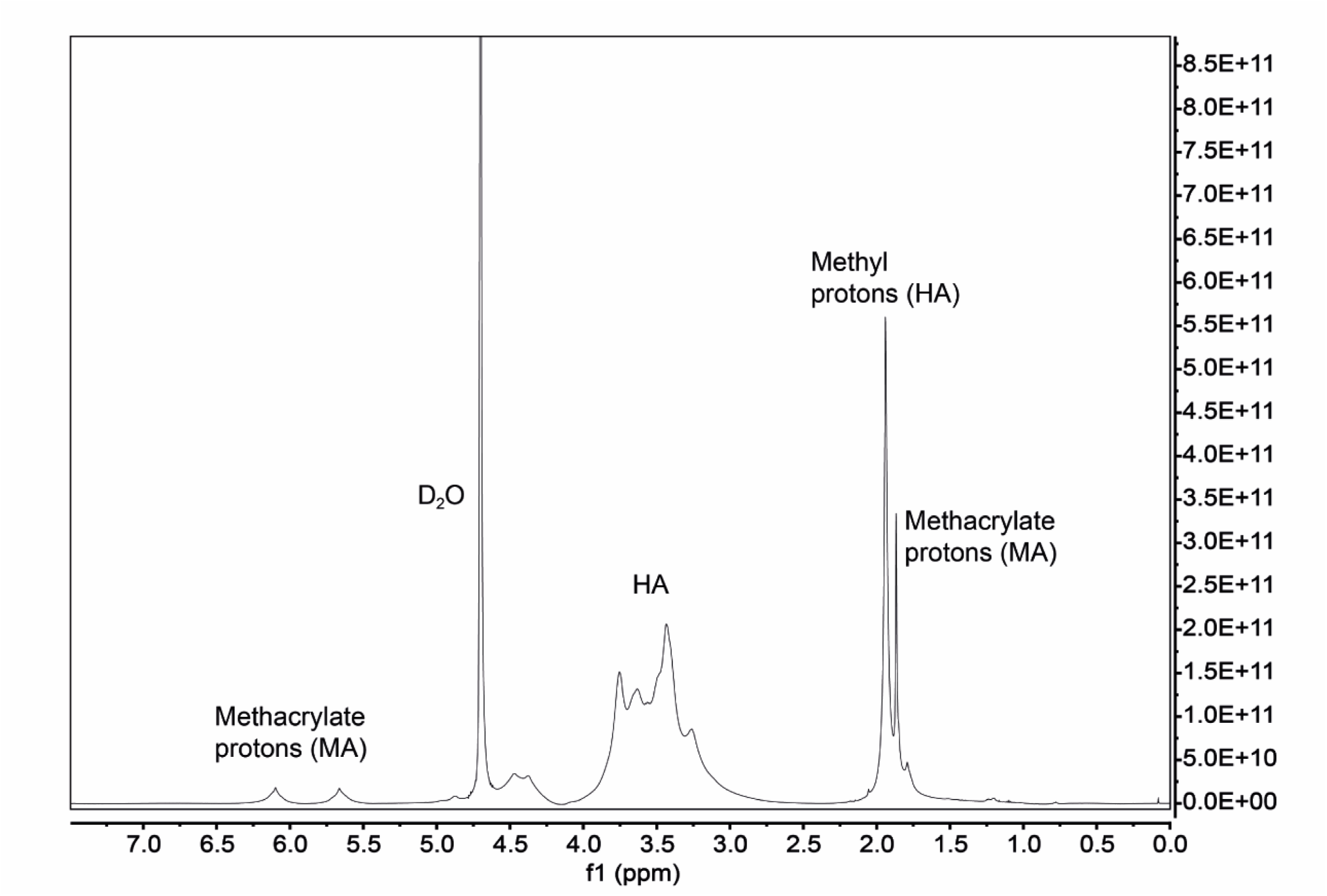
NMR spectra of synthesized HA-MA

**Sup. 2.**
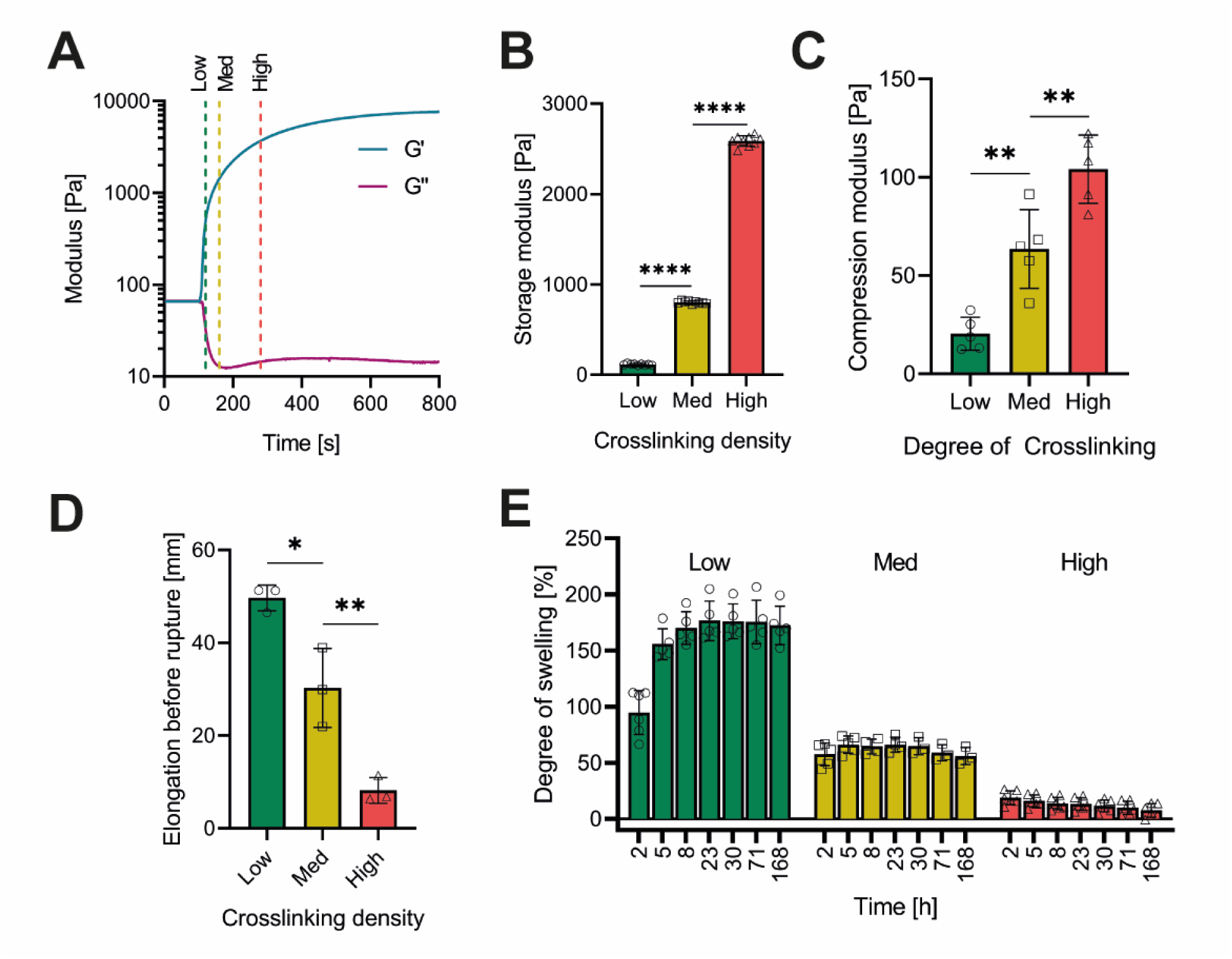
Mechanical properties of HA-MA gels. (**A**) Photocrosslinking behavior of HA-MA with three different time points representing different degrees of crosslinking (*Low*, *Med*, *High*) (**B**) Storage modulus significantly differs between samples (F (_2,24_) = 12164, P<0.001) (**C**) Compression modulus between these samples is significantly different (F (_2,12_) = 34, P<0.001) (**D**) Rupture of a dumbbell shaped samples happens after significant different elongation (F (_2,6_) = 44.24, P<0.001) (**E**) Swelling ratio is different between samples prepared with different crosslinking degrees

**Sup. 3.**
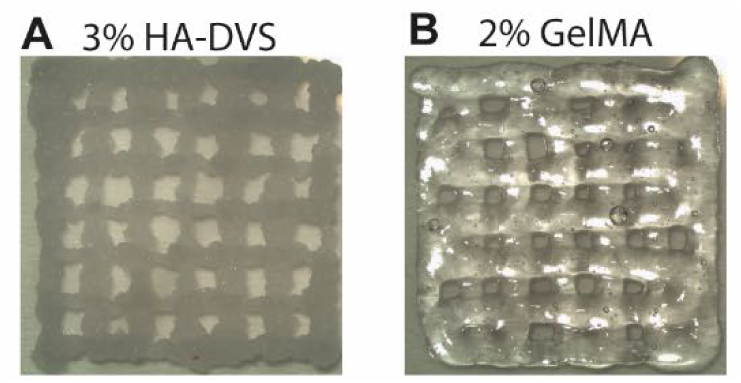
3D printed GelMA and HA-DVS. entangled microstrands prepared and printed from HA-DVS (**A**) and GelMA (**B**)

**Table S1.** Mechanical Properties of Different HA-MA Bulk Gels

|  | Crosslinking<br>time | Storage<br>modulus | Compression<br>modulus | Maximum<br>elongation |
| --- | --- | --- | --- | --- |
| Low | 20 s | 114.1±8.1 Pa | 20.4±7.5 Pa | 49.7±2.3 mm |
| Med | 60 s | 803.6±14.1 Pa | 63.6±17.9 Pa | 30.3±6.9 mm |
| High | 180 s | 2587.8±54.3 Pa | 104.2±15.6 Pa | 8.2±2.3 mm |

**Movie 1: Porosity of entangled microstrands**

